# The dual interpretation of edge time series: Time-varying connectivity versus statistical interaction

**DOI:** 10.1101/2024.08.29.609259

**Authors:** Haily Merritt, Amanda Mejia, Richard Betzel

## Abstract

Functional connectivity (FC) is frequently operationalized as a correlation. Many studies have examined changes in correlation networks across time, claiming to link time-varying fluctuations to ongoing mental operations and physiological processes. Other studies, however, have called these results into question, noting that statistically indistinguishable patterns of time-varying fluctuations can be obtained by windowing synthetic time series generated from ground-truth stationary correlation structure. Recently, we developed a technique for tracking rapid (framewise) fluctuations in network connectivity over time. Here, we show that these “edge time series” are mathematically equivalent to interaction terms in a specific family of general linear models. We exploit this fact to further demonstrate that time-varying connectivity carries explanatory power above and beyond brain activations. This observation suggests that time-varying connectivity is likely more than a statistical artifact.

**SUMMARY:** Brain activity and connectivity have been linked to ongoing behavior and mentation but usually in isolation and almost never in the same model. Here, we show that “edge time series” – a recently proposed method for tracking moment-to-moment connectivity changes – are equivalent to an interaction term in a linear model. By including terms for activations in the same model, it provides an elegant framework for assessing the relative explanatory power of edges and activations. In our work, we use this modeling framework to study time-varying behavior in zebrafish, worms, and humans. We find that connectivity contains unique explanatory power above and beyond activity.

## INTRODUCTION

Functional connectivity (FC) refers to statistical dependencies between activity recorded from distinct neural sources–e.g. cells, neuronal populations, brain areas [1, 2]. Frequently, FC is operationalized as the bivariate product-moment correlation – highly correlated neural elements are considered to be functionally connected [3]. This is especially true at the large scale [4], where whole-brain correlation networks are estimated from functional magnetic resonance imaging blood-oxygen level-dependent (fMRI BOLD) data and used to study the brain’s functional organization [5–7], and its links with cognition [8], disease [9], and development [10].

Most studies of FC estimate a single “static” connectivity matrix whose weights represent the mean coupling strength between brain regions over an entire scan session (usually on the order of 5-30 minutes). Others have argued that there may be benefit to examining time-varying fluctuations in correlation structure over shorter timescales [11–13], linking these changes with network measures [14–17], dynamic fluctuations in arousal [18], and consciousness [19] (among many others).

Time-varying FC (tvFC) is often estimated using a windowing procedure, wherein time series are segmented into (possibly overlapping) windows and FC estimated independently for each window using only those observations that fall within that window [20– 22]. Though popular, this technique has a number of drawbacks. First, the windowing procedure leads to temporal imprecision; correlations estimated in any given window cannot be uniquely attributed to any one time point. This induces “blurring” or “smoothing” that effectively prohibits the detection of rapid changes in coupling structure. Second, a number of studies have suggested that the observed tvFC has artifactual origins [23–26]. These studies generate synthetic time series from ground-truth stationary correlation structure. They show that, when windowed, these time series exhibit temporal variability with statistical properties that are largely consistent with what we observe when the same windowing techniques are applied to empirical time series data.

Recently, we proposed a framework for addressing the first limitation [27]. Specifically, we showed that, for correlation networks, the correlation between nodes *i* and *j* can be decomposed into its time-varying contributions without any windowing [28–30] (see Figure 1 for a schematic illustration of this decomposition). Briefly, this decomposition involves first calculating the element-wise product between pairs of z-scored time courses. While the familiar correlation coefficient is the mean value of this time course, we obtain an estimate of instantaneous co-fluctuations – the edge weight between nodes *i* and *j* – by simply omitting the averaging step. A growing number of studies have examined these “edge time series” in greater detail, exposing whole-brain co-fluctuation events [31–34] and links to phenotypic measures [35, 36], including hormone fluctuations [37], naturalistic stimuli, [38], attention, [39], surprise [40], and the subject-specificity of FC [41, 42].

**Figure 1.**
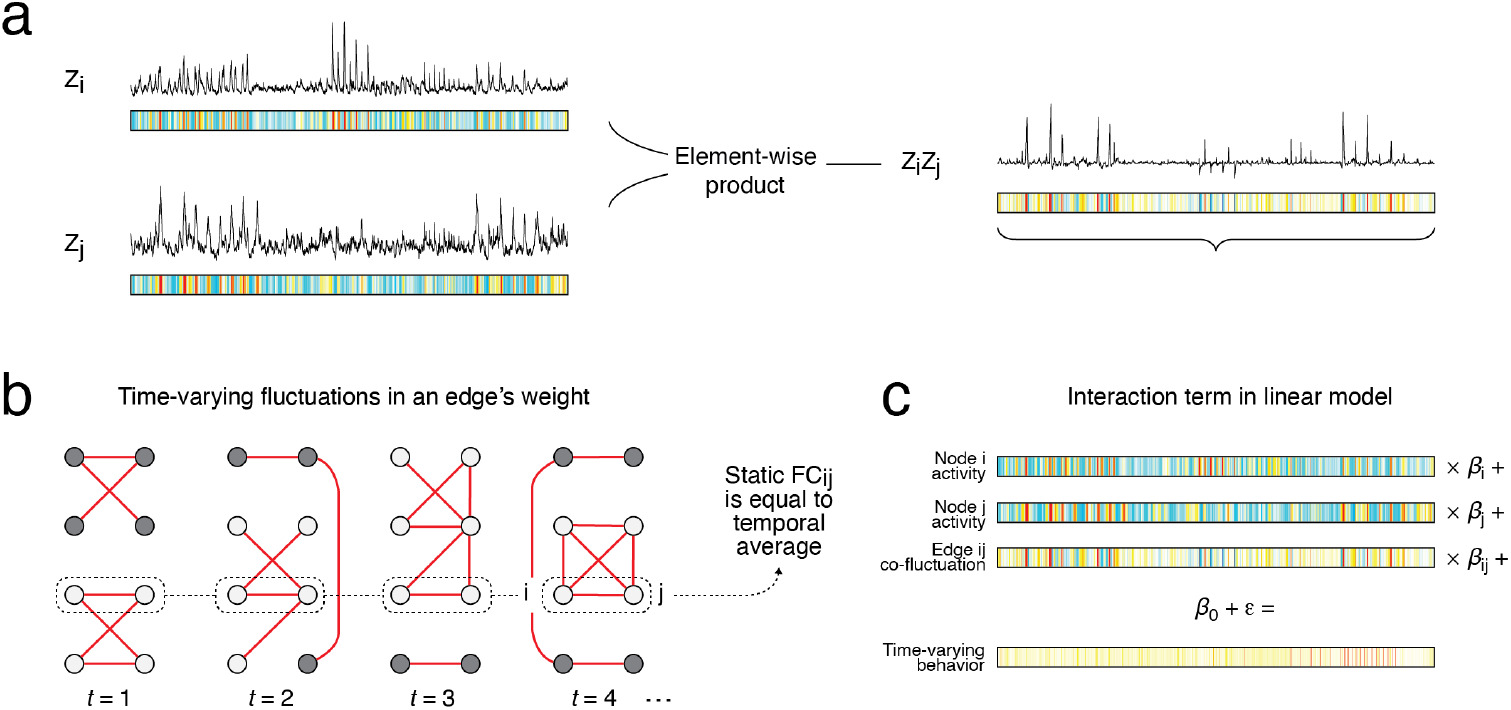
Schematic illustration of edge time series derivation and interpretation. (*a*) Functional connectivity is typically defined as a correlation. Given two z-scored time series of length *T*, **z**_*i*_ and **z**_*j*_, we can calculate their correlation as 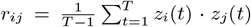. As an intermediate step in this calculation, one creates the co-fluctuation time series, **r**_*ij*_ = [*r*_*ij*_ (1), …, *r*_*ij*_ (*t*)] = [*z*_*i*_(1) · *z*_*j*_ (1), …, *z*_*i*_(*T*) · *z*_*j*_ (*T*)]. (*b*) In most applications, this time series is interpreted as an instantaneous estimate of the weight between nodes *i* and *j*. That is, it serves as a estimate of edge’s weight across time. (*c*) An alternative interpretation of co-fluctuation time series is as the interaction term in a specific class of linear models. Specifically, if we want to explain some time-varying measure of behavior, physiology, or cognition, then we can construct a multi-linear model with regressors corresponding to the z-scored activity of nodes *i* and *j*, as well as their interaction term. This interaction term is mathematically equivalent to the edge time series. Thus, in this second view, edge time series are not directly interpreted as a measure of connectivity, but as a statistical construct.

Here, we make two new contributions. First, we show that edge time series are equivalent to the interaction term in a classical linear regression analysis. This observation is important, as our original rationale for studying edge time series was entirely practical; edge time series were an easily calculated substrate from which we could construct “edge-centric” networks [29, 43–46] while still conforming with standards in human imaging and network neuroscience. Second, because edge time series can be recast as the interaction term in multilinear models, we can easily construct models whose regressors correspond to both activations and connections, allowing us to assess their relative contributions in explaining time-varying behavior. By including these terms in the same model, activations and connections compete for the same variance. If the interaction term is statistically significant, then we can conclude that timevarying connectivity carries explanatory power above and beyond that of activity alone. This observation would suggest that the origins of time-varying fluctuations in connectivity is not simply a statistical artifact.

Here, we use the above-described framework to study fictive swimming and eye movements of larval zebrafish [47], the mating behavior of the nematode, *C. elegans* [48], and responses to dynamic, naturalistic stimuli (movies) in humans [49]. Across phylogeny, scales, and recording modalities, we report evidence that edge time series contribute explanatory power beyond that of the activations. These findings build on other recent efforts to use edge time series for explaining dynamic fluctuations in attention [50] and surprise [40] and call into question the longstanding criticism of time-varying connectivity as purely stochastic and artifactual [23– [26, [51]. Finally, our work shares conceptual links with other foundational approaches for linking time-varying measures of behavior with recordings of brain activity, e.g. psychophysiological interactions (PPI) [52, 53], positioning edge time series as a potent framework for studying brain network organization, its reconfiguration over time, and its behavioral correlates.

## RESULTS

### Edge time series are mathematically equivalent to the interaction term in linear models

Though originally introduced as an intermediate step in the construction of higher-order edge-centric networks [29, 45, 54], edge time series are more often used to study tvFC [27]. To construct the edge time series for the node pair, {*i, j*}, we calculate the element-wise product of their z-scored activity time courses. That is: **r**_*ij*_ = **z**_*i*_ · **z**_*j*_, where **z**_*i*_ = [*z*_*i*_(0), …, *z*_*i*_(*T*)] and · denotes the element-wise product.

Edge time series have a straightforward interpretation; the value of *r*_*ij*_(*t*) corresponds to the instantaneous co-fluctuation between nodes *i* and *j*. The sign of *r*_*ij*_(*t*) is positive when their activity at time *t* is deflecting in the same direction relative to their respective means. The amplitude of *r*_*ij*_(*t*) captures how far above or below their means each time course is.

Historically, edge time series have been treated as estimates of instantaneous connectivity. They have been used to discover high-amplitude “events” [28, 31, 32, 37, 55], shed light on the origins of modular brain systems [56], examine associations with cognition and attention[39], and study individual differences and fingerprinting [35, 41, 57].

However, we assert that edge time series have an alternative (but not mutually exclusive) interpretation when viewed through the lens of elementary statistics. Suppose we wanted to explain some time-varying behavior, **y** = [*y*(1), …, *y*(*T*)], with brain imaging data. Arguably the simplest approach is to explain this behavior in terms of activations (ignoring, for the time being, hemodynamic lags and serial correlations). We would write this model as:

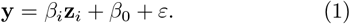

As before, we let the variable **z**_*i*_ correspond to an activity time series–e.g. the fMRI BOLD time series recorded from a region *i*. For the sake of concreteness, one might imagine using this model to explain fluctuations in pupil diameter (an indirect measure of arousal [58]) in terms of activity recorded from a voxel/grayordinate. In this example, **y** and **z**_*i*_ correspond to the pupil diameter and fMRI BOLD time courses, respectively.

However, we could create a more complex model that includes additional predictors, including the activity of another brain region, *j*. We might, then, also include the interaction term associated with the activity of regions *j* and *j*. In that case, we would write our model as:

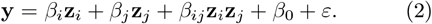

Note here that the interaction term is calculated as the element-wise product of the two activity time courses, **z**_*i*_ and **z**_*j*_. Because the activity time series are expressed as z-scores, this product is mathematically equivalent to our calculation of instantaneous cofluctuations and therefore the edge time series. Thus, elementary statistics facilitates a convenient interpretation of edge time series as the interaction of two z-scored regressors.

This equivalence confers several useful benefits. Beyond expanding the usefulness of a widely used tool (the linear model) in behavioral neuroscience, this framework allows activation and connectivity (in the form of edges) to “compete” for the same variance in the behavioral measure, **y**. We explore implications in the following sections.

### Edges explain time-varying behavior above and beyond activity

One of the longstanding assumptions in network neuroscience is that time-varying fluctuations in the coupling of neural elements to one another relate to ongoing behavior. This assumption has not gone unchallenged; models of synthetic fMRI BOLD data exhibit patterns of tvFC similar to those of empirical data, despite stationary correlation structure and no cognitive or physiological component in their generative model. These simulated results suggest that observed tvFC may largely reflect sampling variability as opposed to fluctuations in the support of behavior.

We reason, however, that if edge time series can explain time-varying behavior beyond what can be explained by activity alone (node time series), then timevarying functional connectivity (as operationalized by an edge time series) is unlikely a simple statistical artifact. Here, we directly examine this hypothesis by fitting multi-linear models to behavioral data acquired alongside three brain imaging datasets: seven timevarying behaviors of larval zebrafish (five measures of fictive swimming and two measures of eye movement), annotations describing the semantic content of naturalistic stimuli (human fMRI), and five measures associated with *C. elegans* mating behavior. We show that, despite variability in imaging modality, phylogeny, and spatial scale, many edges carry explanatory power. In this section we describe results from the zebrafish dataset; we report human and worm results in subsequent sections. See Fig. S1 for a brief overview of the zebrafish dataset and summary network statistics.

An important note. In the following sections, all models were fit at the level of individuals. The results of those individual-level models were then aggregated to generate group-level summaries. We note that this approach is at odds with the convention in non-invasive imaging to perform statistical tests at the group-level (here, this would mean estimating a single set of regression coefficients for the entire cohort). Our rationale for fitting individualized models is multi-fold. First, our datasets all included large amounts of data per individual (fish, worm, person). We are therefore more likely to be powered to detect effects at the individual level than, say, a study in which a small amount of data was collected for each individual. Second, there is a renewed emphasis in non-invasive neuroimaging to focus on individualized features, including responses to stimuli and tasks. Our approach is consistent with this idea. Further, by generating group-level summaries of parameters estimated at the individual level, we obtain a clearer picture of where there exists consensus (or dissensus) across individuals.

In greater detail, we fit multi-linear models at each edge, {*i, j*. The model included regressors **z**_*i*_, **z**_*j*_, and **z**_*ij*_, which represented the z-scored activity of nodes *i* and *j* and their interaction term–i.e. the edge time series. We fit two versions of these models: In the main text we describe models fit to pre-whitened data – a procedure used to reduce the inflated effect sizes that arise from serially correlated time series. In the supplementary material we report results using a version of the same data that *does not* include pre-whitening as processing step. The results of both models largely converge in that both identify a plurality of statistically significant interactions (see Fig. S2). In the main text, we focus on the pre-whitened data.

The fitted model yields several useful terms: the *β*_*i*_ and *β*_*j*_ regression coefficients (Fig. 2)c, left and center), which describe the contributions of brain activity to the model; *R*^2^, which describes the variance explained by the model (Fig. 2)b); and the interaction coefficient, *β*_*ij*_, which describes the contribution of edges to the model performance (Fig. 2)c, right). Our focus is on the interaction coefficients for all edges, {*i, j*}, which we store as an [*N* × *N* ] matrix for each behavioral measure (Fig. 2a). We refer to this matrix as the “interaction matrix.”

**Figure 2.**
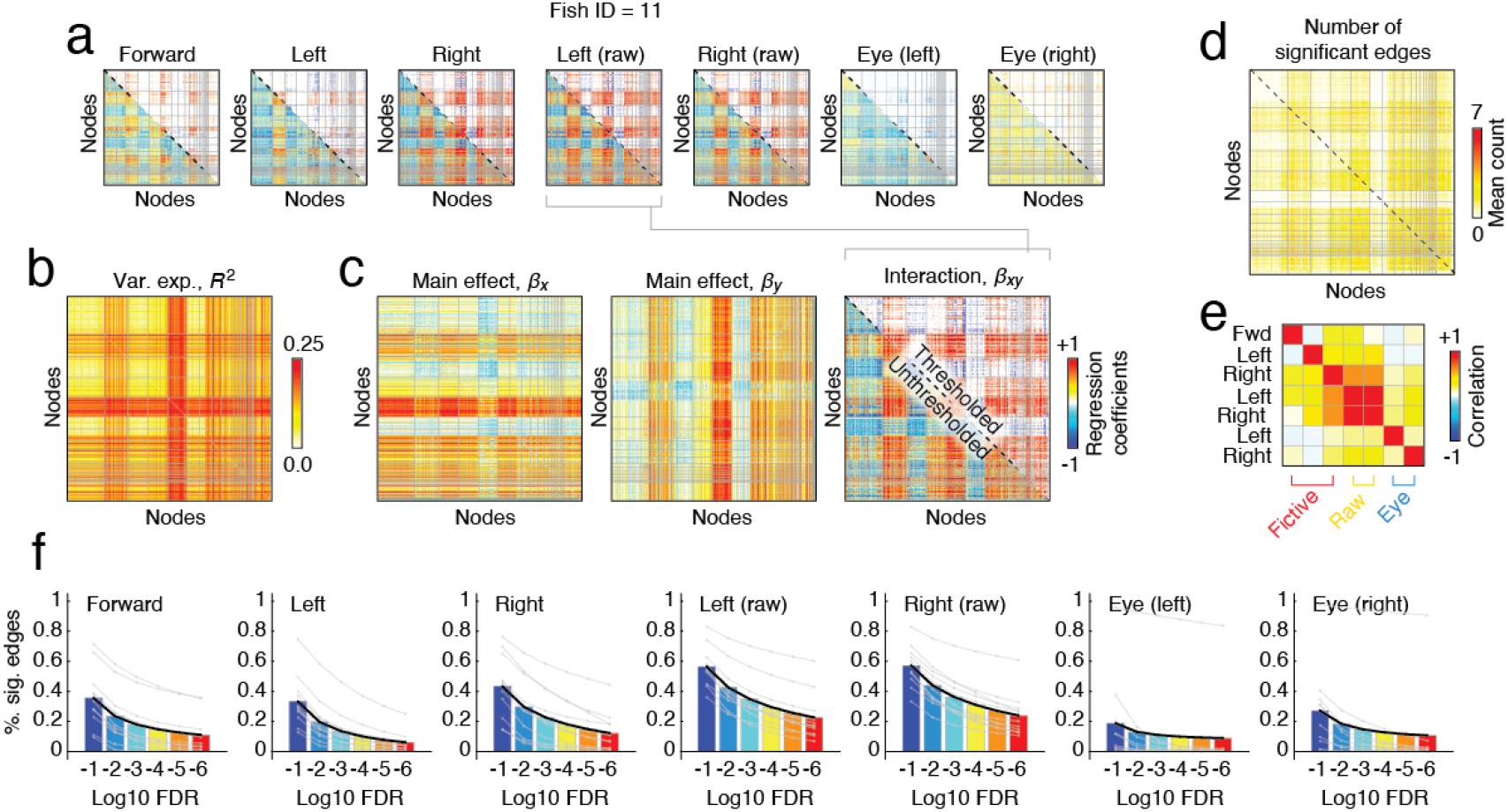
Regression analysis of edge time series. Here, we show representative results from a single fish (Fish ID 11). (*a*) Matrices of regression coefficients, *β*_*xy*_. In general, these matrices are symmetric; the lower triangle shows unthresholded coefficients while the upper triangle shows the same coefficients after statistical testing (false discovery rate fixed at *q* = 0.01). We focus further on the matrix for leftward movements (fictive). (*b*) Variance explained, *R*^2^, by each model. (*c*) Regression coefficients for the activity of regions *x* and *y* and the interaction term, {*x, y*}. We also identify consistently significant edges at the group level. (*d*) The number of behaviors for which a given edge was statistically significant at the group level. (*e*) Spatial similarity (correlation) of whole-brain regression coefficients. Most importantly, we calculated, at varying critical values and separately for each of the seven behavioral measures, the fraction of edges that meet the criteria for statistically significance. In panel *f*, we show the results of this analysis for each measure (sub-panel) and where we fix the false discovery rate at 10^−1^, 10^−2^, 10^−3^, 10^−4^, 10^−5^, and 10^−6^. The black line in each subplot represents the population mean. Gray lines represent individual animals.

We investigated the interaction terms in detail. First, we asked simply whether the any of the interactions were statistically significant–i.e. whether edges contributed explanatory power above and beyond nodal activations. To do this, we employed a mass-univariate testing procedure wherein we tested the hypothesis that the interaction term was statistically significant at each edge. To control for multiple comparisons (we repeated this testing procedure for *N* (*N* − 1)*/*2 node pairs), we adjusted the critical *p*-value to accommodate a small fraction of false positives. We tested false discovery rates (FDR) ranging from permissive to conservative spanning the interval 10^−1^ to 10^−6^. In general, we found robust evidence that at least some edges always contributed explanatory power spanning the range of FDRs considered here (Fig. 2f). At *q* = 10^−1^ = 0.1, the fraction of significant edges/interactions ranged from ≈40%-70% across behaviors. With more conservative control (*q* = 10^−6^), the fraction of significant edges remains substantial, ranging from ≈10%-40% across behaviors.

Next, we tested the behavioral specificity of the significant edges. To do this, we calculated the population mean regression coefficient for each edge, {*i, j*}, and each behavior. Then, assuming equal degrees of freedom across fish, we calculated the *p*-value at each edge under the *t*-distribution and corrected for multiple comparisons using the Benjamini-Hochberg approach [59]. For a given behavior, this yielded a 0 (not significant) or 1 (significant) value at each edge. We averaged these values across behaviors, yielding a single *N* × *N* map of edge significance frequency (Fig. 2d). The most consistent edges linked neurons in the rhombencephalon and diencephalon (Fig. S3). Nonetheless, the whole-brain interaction patterns were heterogeneous; although raw left and right turns were highly similar, the remaining measures were not strongly correlated with one another (Fig. 2e). This general pattern holds when, instead of group-level parcels (nodes), we use subject-specific and functionally-defined parcels (see Fig. S8).

As a final methodological note, we point out that throughout the above analyses we do not convolve the behavioral data with a kernel to model the slow kinetics associated with calcium fluorescence. In the supplementary material, we explore the effect of convolution and observe that, in general, results not only hold, but that the explanatory power of models is enhanced and the number of statistically significant edges increased when the behavioral data is convolved with the appropriate kernel (Fig. S4). Thus, the results reported here in the main text are likely overly conservative.

In summary, these observations suggest that timevarying connectivity contains signal and not merely noise. More than that, connectivity explained timevarying behavior above and beyond activity. This finding holds across a range of false discovery rates, behaviors, imaging modalities, phylogenies, and spatial scales. Having built confidence in the predictive power of edge time series, we next examine how the association between edge time series and time-varying behavior varies with context.

### Interaction patterns are sensitive to context

In the previous section we showed that, across a range of behaviors, edges carry explanatory power above and beyond brain activity recordings. We characterized the contributions of edges to explanations of behavior in terms of whole-brain interaction matrices. How malleable are these interaction patterns? Are they preserved across experiments (different conditions) or are they flexible and capable of reconfiguration?

To test this, we segmented the entire recording session into distinct stimulus segments. Because stimulus conditions repeated within a given recording session, blocks were discontinuous (see Fig. 3a). We then estimated interaction matrices separately for each stimulus, combining across discontinuous blocks (Fig. 3b and c), and calculated the pairwise similarity (correlation) between all interaction matrices from all conditions (Fig. 3d). This approach allowed us to classify pairs of interaction patterns according to two different properties: a given pair of interaction matrices could correspond to the same *versus* different stimulus or they could correspond to the same *versus* different behavior (see Fig. 3e,f). For each property, we tested whether same-property pairs were more similar than different-property pairs. While interaction patterns were statistically indistinguishable when comparing the same and different behaviors, we found that interaction patterns estimated within a given stimulus condition were more similar to each other than to patterns between different conditions (paired sample *t*-test; *p* = 4.4 × 10^−6^).

**Figure 3.**
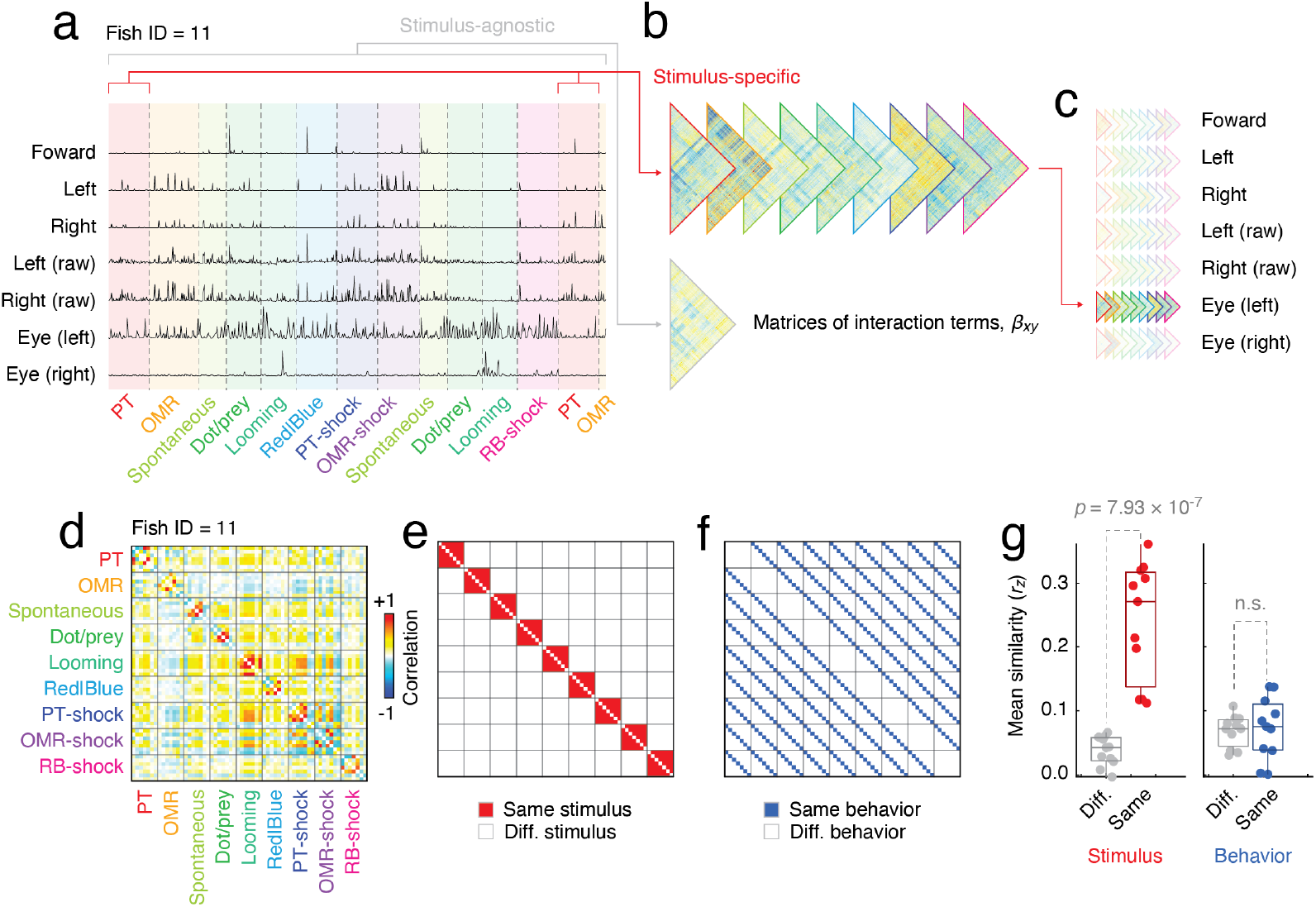
Stimulus-specificity of whole-brain regression coefficient maps. In our previous analyses, we estimated interaction coefficients for each behavioral variable using data from the entire recording session. However, each recording was composed of multiple stimulus conditions (plus a non-stimulus “spontaneous” condition). Here we show results after estimating regression coefficients and separating time points by condition. Panels *a*-*c* illustrate the general methodological approach. For each of the seven behaviors, we estimate condition-/stimulus-specific interaction terms. (*d*) Spatial similarity of vectorized whole-brain interaction matrices. We tested two hypotheses. Are interaction patterns stable within conditions but variable across conditions? If so, we expect the similarity to cluster around stimulus/condition (*e*). On the other hand, are the interaction patterns stable across conditions so that irrespective of condition/stimulus, the interaction patterns are similar for the same behavior (*f*). (*g*) We find that interaction patterns are more similar within stimulus/condition than between, but no such effect for behavior.

These observations suggest that although we can identify meaningfully predictive edges for a variety of behaviors, precisely *which* edges are statistically significant depends on the context. Specifically, significant edges are more similar under the same stimulus than a different stimulus, but the same is not true for the same behaviors. Next, we extend these results across phylogeny, spatial scales, and imaging modalities.

### Edges explain behaviors of different phylogeny, at different spatial scales, and different imaging modalities

In the previous sub-sections, we showed that edge time series are equivalent to interaction terms, that edges carry explanatory power above and beyond activations, and that their contributions to explanations are context specific. Here, we extend and corroborate these observations, reporting similar effects using two additional datasets: human functional MRI during movie-watching and *C. elegans* light-sheet microscopy during mating behavior.

We analyzed human imaging data recorded while participants watched four movies (sequences of movie clips interspersed by black screens; approximately 15 minutes long) [49]. The movies were accompanied by rich annotations of their semantic content as framewise indicator variables. We used a clustering algorithm to group these vectors into collections of terms that cooccurred (see Fig. 4a-d) and, as in the previous section, fit multilinear models to explain these groups in terms of activity (nodes) and interactions (edges).

**Figure 4.**
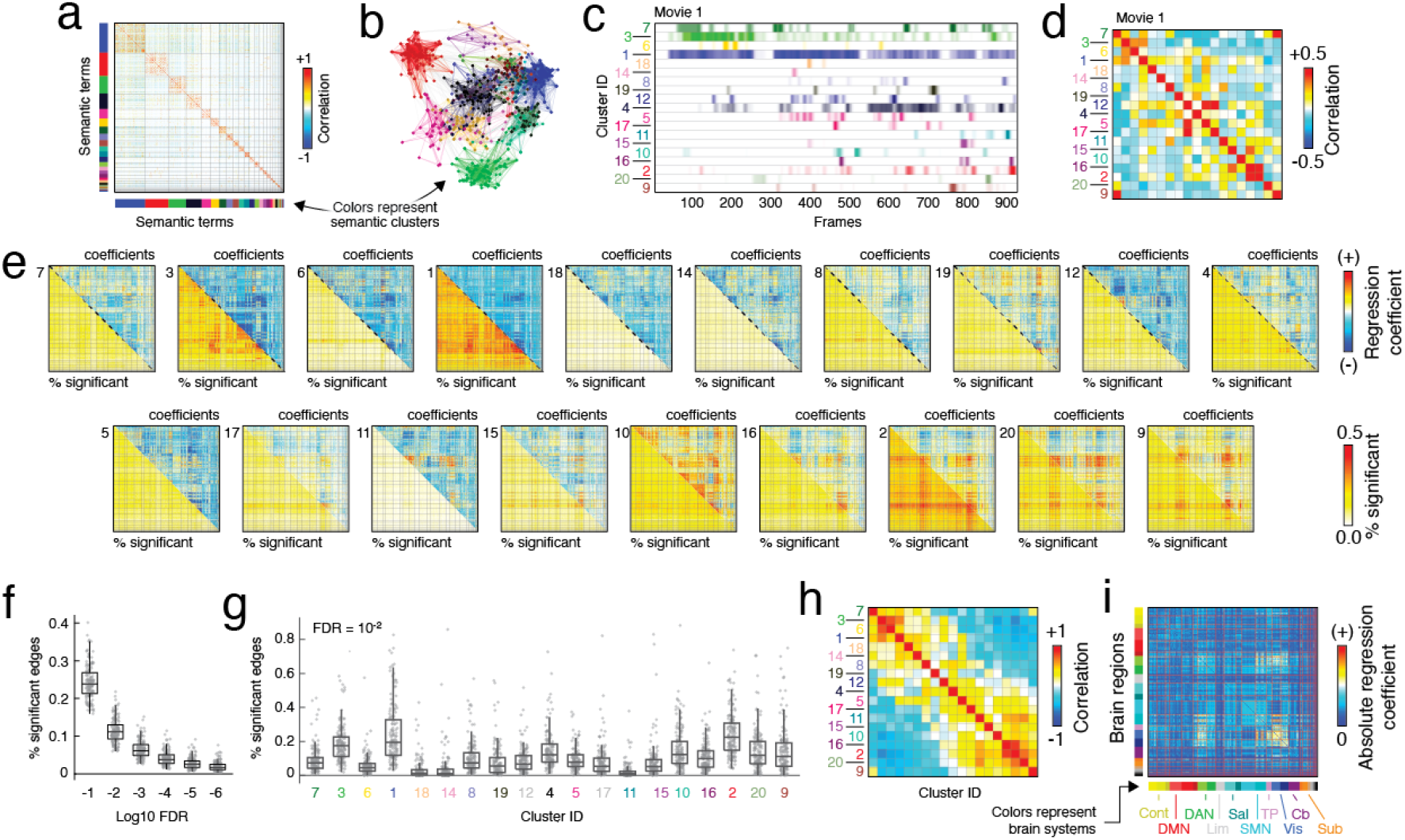
Analysis of human movie-watching data. (*a*) Correlation matrix of 859 terms/annotations across four movies; rows and columns sorted by community structure. (*b*) Force-directed layout for communities shown in *a*. (*c*) Time courses for each cluster: mean annotation time course for all terms assigned to a given cluster and subsequently convolved with hemodynamic response function (Movie 1). (*d*) Correlation matrix of cluster time courses. (*e*) Interaction matrices for 19/20 clusters for Movie 1. The upper triangle shows the subject-averaged interaction coefficients; the lower triangle displays the fraction of participants for which a given edge was statistically significant. (*f*) Fraction of edges that pass statistical significant at different levels of false-discovery corrections. (*g*) For an allowed false discovery rate of *q* = 0.01, the fraction of significant edges calculated separately for each cluster. In panels *f* and *g*, points correspond to participants. (*h*) Similarity (correlation) of unthresholded interaction matrices. (*i*) For each edge, we calculated the mean absolute regression coefficient averaged across all annotation clusters. Note that here we show results from Movie 1 in which 19/20 clusters are present (no terms assigned cluster 12 appear in this movie and so we omit that cluster in our plots).

As in the previous section, we found ample evidence that co-fluctuations captured by edges carry information about a time-varying stimulus. Edges explain timevarying behavior above and beyond the level of activity (see Fig. 4e for regression coefficients and the fraction of participants for which a given coefficient was statistically significant). Importantly, although the fraction of significant edges decreases as our correction for multiple-comparisons becomes more stringent, we nonetheless find evidence that edge time series meaningfully explain time-varying behavior across the full range of corrections tested here (Fig. 4f). Interestingly, there was considerable heterogeneity across annotation groups. For instance, the average fraction of significant edges associated with cluster 1 was around 20%, whereas cluster 11 was close to zero. There was also marked heterogeneity in the similarity of interaction matrices (Fig. 4h). Nonetheless, there was convergence in terms of which edges were statistically significant (proportional to their regression coefficient amplitude), with the largest coefficients converging on sensorimotor networks include visual, dorsal attention, and somato-motor systems (Fig. 4i).

In addition to human imaging data, we also fit multi-linear models to neuron-resolution recordings from *C. elegans* during mating behavior [48] (see Fig. 5a-e). Specifically, we considered behavioral variables “Velocity”, “Tail Curvature”, “Distance to tips”, “Distance to vulva”, and “Spicule protraction” (see **Materials and Methods**). Further, the mating behavior could be broken down into distinct periods, allowing us to assess to what extent interactions matrices are context-dependent.

**Figure 5.**
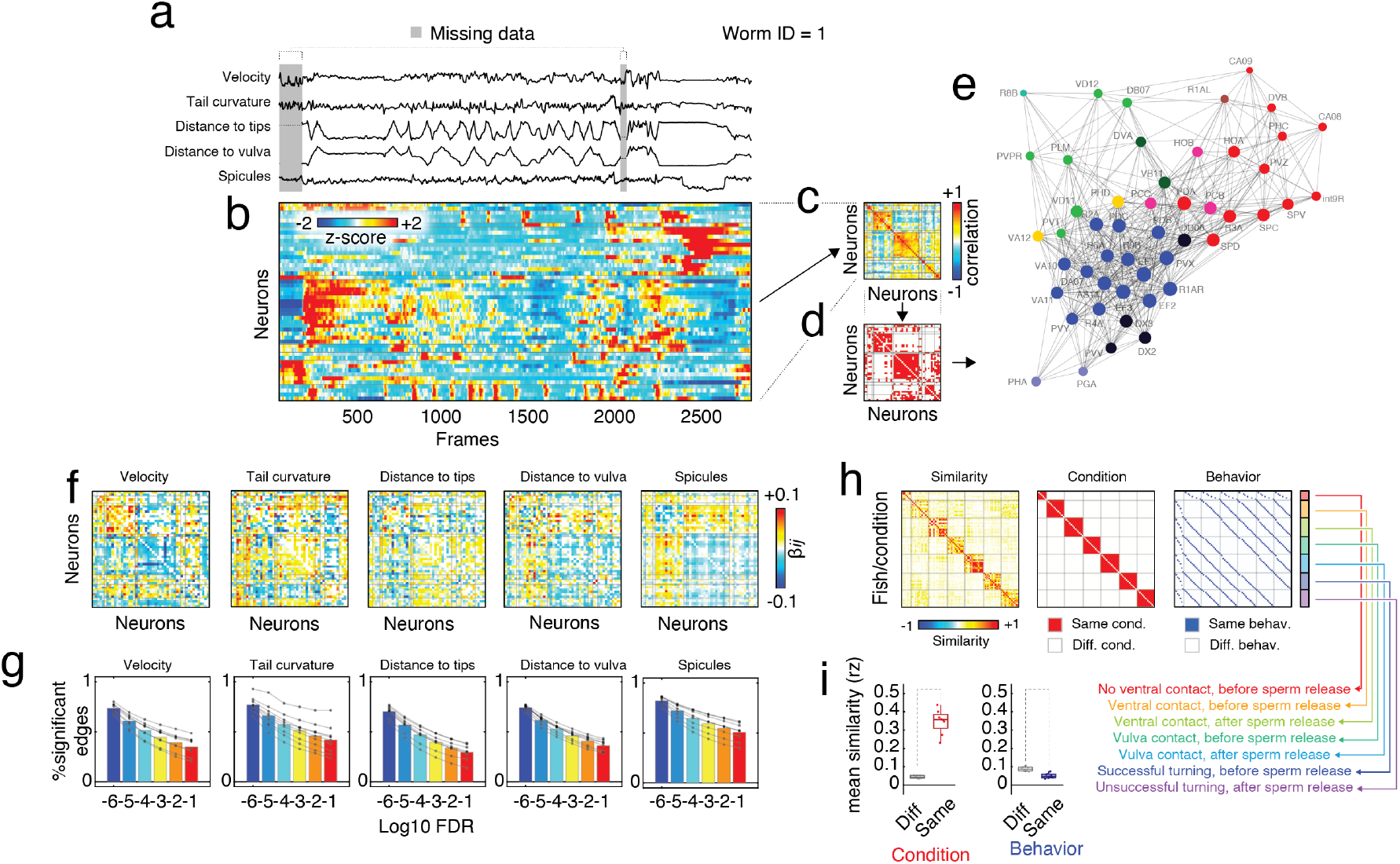
Analysis of *C. elegans* data. (*a*) Time-varying measures of behavior for representative worm. (*b*) Z-scored activity from individual neurons. The correlation structure, thresholded correlation matrix, and spring-embedded matrix are shown in panels *c*-*e*, respectively. As with zebrafish and human imaging data, we used these time series to model time-varying behavior. In this case, we focus on five behavioral measures associated with mating. (*f*) Interaction matrices for five measures. Note that panels *a*-*f* come from a single representative worm (ID = 1). In panel *g*, we pool across different worms and plot the fraction of edges that exhibit statistically significant interactions while controlling the false discovery rate at different levels. We also investigated the condition-specificity of the interaction patterns. (*h* and *i*) Like zebrafish, we find that interaction patterns are stable (similar) within condition.

In a near-perfect recapitulation of our findings with zebrafish, we observed statistically significant interactions for every worm and for every behavior (Fig. 5f,g). As with the zebrafish, the interaction patterns were more stable within conditions compared to between conditions (Fig. 5h,i). Moreover, these results persisted across a range of significance thresholds (Fig. 5g).

In summary, our findings held across phylogeny, spatial scales, and imaging modalities. The stability of these results further strengthens the claim that edge time series meaningfully predict time-varying behavior and that time-varying functional connectivity defined this way is behaviorally relevant. As a final observation, we note that the whole-brain interaction matrices were also broadly related to time-averaged correlation matrices–i.e. FC (Fig. S9). These observations further ground our approach in familiar constructs and present opportunities for future work to investigate these relationships in greater detail.

## DISCUSSION

Here, we make two broad claims. First, we show that edge time series are equivalent to the interaction term in a specific multilinear model. These models allow us to directly test whether time-varying behavior is best explained by fluctuations in activity or connectivity (or both). Second, we apply these models to three diverse datasets that span imaging modalities – lightsheet microscopy and functional MRI – and phylogeny – fish, humans, and worms – and show that, for a wide range of behaviors, time-varying connectivity helps to improve explanatory power. Our results suggest that time-varying connectivity is more than a statistical artifact and that the coordination of neural elements across time is linked to ongoing behavior.

### Edge time series *are* interaction terms

Although independently discovered by van Oort *et al*. [60], edge time series have garnered interest of late as an approach for estimating tvFC. They have some advantages over sliding-window methods, in that they estimate framewise coupling and therefore offer greater temporal precision. Further, the temporal average of edge time series is exactly the bivariate product-moment correlation coefficient, positioning edge time series as a means of unwrapping FC to examine how single frames contribute to the average coupling between two neural elements. On the other hand, co-fluctuation amplitudes are unbounded, which occasionally makes their interpretation challenging. Further, the derivation of edge time series in Faskowitz *et al*. [29] was motivated by mere practicality; edge time series were simply an intermediate step in estimating edge-centric FC [45, 54].

Here, we show that edge time series are equivalent to the interaction term in linear models, situating them on the *terra firma* of multilinear models and elementary statistics. Although previous studies have used edge time series as regressors in linear models to explain behavior [39, 40], the mathematical parallel between edge time series and interaction terms has, to our knowledge, not been noted elsewhere.

This equivalence also presents an interesting lens through which we can interpret static FC. Interaction terms are included in a linear model when it is hypothesized that the relationship between a regressor and a response variable is dependent upon the value of another regressor. It is typically modeled as the product of the two regressors. It is also well-established that the temporal average of an edge time series is exactly the bivariate product-moment correlation [30, 61]. Therefore, we can interpret FC as the mean interaction between two time series.

### Connectivity and activity are needed to explain behavioral fluctuations

tvFC has proven a complicated topic within network neuroscience [11]. Many groups have presented compelling evidence that the coupling structure (connectivity) of brain activity is not fixed over time [12, 14, 16, 38, 62, 63] and that these fluctuations may be linked to ongoing behavioral fluctuations [64, 65]. Concurrently, other groups have argued the opposite, noting that observed fluctuations in FC are consistent with what might be predicted by sampling variability [23–25, 51].

Here, we use tvFC estimated at framewise resolution to explain a wide range of time-varying behaviors, ranging from fictive movement to movie content to mating behavior. Importantly, and in contrast with most previous studies (though see Benisty *et al*. [64] for a notable exception), we allow time-varying connectivity estimates to compete with activation time courses in explaining behavior. We find robust statistical evidence supporting the hypothesis that edge time series offer unique explanatory power beyond that of activations. In fact, our mass-univariate approach for model-fitting/-testing may be underpowered for detecting significant effects; future studies may be able to leverage multi-variate modeling approaches to improve statistical power [66].

Notably, we found that interaction patterns are sensitive to context; brain regions are not necessarily coordinating in the same way each time the same behavior is performed. Other work has emphasized that associations between behavior and connectivity can vary across contexts [67]. Together, this suggests that simply identifying brain-behavior associations is only part of the story and that clarifying variation across context– whether temporal or otherwise–supports a richer understanding of the neural underpinnings of behavior.

Although these observations suggest that statistically significant interactions are abundant (and therefore connectivity has some unique explanatory power), they offer no clear mechanism by which connectivity produces or engenders behavior [68]. Nonetheless, these observations can help guide future experimental work. For instance, chemo-/opto-genetic control could be leveraged to disrupt the coordination (interaction) between populations of neurons in an effort to disregulate behavior [69]. This would allow us to test the causal contributions of interactions to ongoing behavior.

## Limitations of the study

This study has a number of limitations. Most notably, though it does link time-varying connectivity (in the form of edge time series) with ongoing behavioral fluctuations, the relationship is explanatory and noncausal. Second, though we aimed to sample multiple species and imaging modalities, the temporal resolution of the fMRI BOLD and light-sheet imaging data are comparable. In fact, this was a deliberate decision, as edge time series are most useful when FC is defined as a correlation, which may not be the appropriate measure for assessing FC for fast time series, e.g. EEG or MEG [70–72]. Accordingly, it remains unknown whether similar analyses using more sensitive measures better-suited for detecting changes in network structure on the millisecond time-scale would yield similar results. With the exception of *C. elegans*, where time courses were derived for individual neurons, the zebrafish and human time series corresponded to aggregates: of single cells in the case of zebrafish and grayordinates in the case of human fMRI BOLD data. While aggregating micro-elements into parcels can enhance noisy signals, it necessarily washes out variability across micro-elements. Therefore, it is unclear whether we would obtain similar results had we carried out these analyses on cells or grayordinates. Doing so is, of course, computationally burdensome. Consider the approximately 80,000 neurons identified in each zebrafish dataset. If we were to apply to those data the same methods described here, would require fitting nearly 3.2 billion models ((80000 · 79999)*/*2).

## Conclusions

In conclusion, we show that edge time series and interaction terms are one and the same for a given class of model. We exploit this relationship to test whether edges – connectivity – or activity are more closely related to time-varying behaviors. We find robust evidence that edges offer unique explanatory power. These observations hold across three datasets and suggest that time-varying FC, at least when measured using edge time series, is likely not statistical noise.

## MATERIALS AND METHODS

### Datasets

#### Zebrafish light-sheet imaging data

Activity from a majority of neurons was recorded in larval zebrafish using light-sheet microscopy [73]. Activity was recorded under spontaneous conditions and also while subjects were presented with a series of visual stimuli. As reported in [47], calcium imaging data was recorded at ≈ 2 volumes/second for ≈ 50 minutes (or ≈ 6800 volumes). As in [73], the calcium indicator, GCaMP6f [74], was expressed pan-neuronally and fused to cell nuclei, allowing for the automatic segmented of cells [75]. The result was continuous fluorescence traces for ≈ 80, 000 cells per subject. As noted in [47], this number accounts for the majority of neurons in the brain, excluding extremely ventral areas. For complete details of data acquisition, see [47].

Rather than analyze networks where nodes correspond to individual cells, we focused on networks where the nodes represented clusters of cells, effectively reducing computational burden and facilitating straightfor-ward interpretation. Our algorithm consisted of the following steps. First, we mirrored all neurons across the hemisphere midline and associated each neuron with one of four large-scale anatomical structures: telencephalon, rhombencephalon, mesencephalon, and diencephalon. We then calculated the total percentage of all neurons assigned to each structure; the number of “parcels” associated with each structure was proportional to these percentages. With *k* = 500 single-hemisphere parcels, this translated into 92, 484, 316, and 108 parcels in the telencephalon, rhombencephalon, mesencephalon, and diencephalon, respectively. We then clustered neurons for each structure independently, so that clusters were homogeneous in terms of their anatomical composition. For each cluster, we estimated its spatial centroid and mirrored those centroids into the other hemisphere, yielding *k* = 1000 bilateral cluster centroids. Finally, we assigned individual neurons to their nearest centroid. Using these assignments, we estimated animal- and centroid-specific activity time series as average z-scored time course of all neurons assigned to that cluster. Note that this procedure diverges from what was reported in Betzel [76] in that neuron-to-parcel assignments are constrained by anatomy and we did not perform global signal regression.

The above procedure yields a shared set of network nodes across zebrafish. While these nodes emphasizes group-level spatial features, facilitate cross-fish comparisons, and enables straightforward summaries of the findings, they may nonetheless distort or average over features unique to each fish. To address this concern, we also derived “individualized” parcellations for each fish that respect fish-specific functional and spatial boundaries. Briefly, using data from the entire recording session, we used modularity maximization to partition neurons into non-overlapping communities. We then mirrored neurons into a single hemisphere and, using spatial information, sub-divided each community until the maximum diameter (largest distance between two neurons in the same cluster) was below a fixed threshold. The sub-clusters were then reflected back into their original hemispheres and representative time courses extracted for each sub-cluster (see Fig. S8).

Along with neural activity traces, these data included several measures of fictive behavior. As described in Chen *et al*. [47], two glass pipettes were attached to the skin on both sides of the tail. Electrodes recorded from the axons of multiple motor neurons, providing a readout of fictive motion [77, 78].

#### Human functional MRI data

The Human Connectome Project (HCP) 7T dataset [49] consists of structural magnetic resonance imaging (T1w), resting state functional magnetic resonance imaging (rsfMRI) data, movie watching functional magnetic resonance imaging (mwfMRI) from 184 adult subjects. These subjects are a subset of a larger cohort of approximately 1200 subjects additionally scanned at 3T. Subjects’ 7T fMRI data were collected during four scan sessions over the course of two or three days at the Center for Magnetic Resonance Research at the University of Minnesota. Subjects’ 3T T1w data were collected at Washington University in St. Louis. The study was approved by the Washington University Institutional Review Board and informed consent was obtained from all subjects.

We analyzed MRI data collected from *N*_*s*_ = 129 subjects (77 female, 52 male), after excluding subjects with poor quality data. Upon defining each spike as relative framewise displacement of at least 0.25 mm, we excluded subjects who fulfill at least 1 of the following criteria: greater than 15% of time points spike, average framewise displacement greater than 0.2 mm; contains any spikes larger than 5mm. Following this filter, subjects who contained all four scans were retained. At the time of their first scan, the average subject age was 29.36 ± 3.36 years, with a range from 22 − 36. 70 of these subjects were monozygotic twins, 57 where non-monozygotic twins, and 2 were not twins.

A comprehensive description of the imaging parameters and image preprocessing can be found in [79] and in HCP’s online documentation (https://www.humanconnectome.org/study/hcp-young-adult/document/1200-subjects-data-release).

T1w were collected on a 3T Siemens Connectome Skyra scanner with a 32-channel head coil. Subjects underwent two T1-weighted structural scans, which were averaged for each subject (TR = 2400 ms, TE = 2.14 ms, flip angle = 8^°^, 0.7 mm isotropic voxel resolution). fMRI were collected on a 7T Siemens Magnetom scanner with a 32-channel head coil. All 7T fMRI data was acquired with a gradient-echo planar imaging sequence (TR = 1000 ms, TE = 22.2 ms, flip angle = 45^°^, 1.6 mm isotropic voxel resolution, multi-band factor = 5, image acceleration factor = 2, partial Fourier sample = 7/8, echo spacing = 0.64 ms, bandwidth = 1924 Hz/Px). Four resting state data runs were collected, each lasting 15 minutes (frames = 900), with eyes open and instructions to fixate on a cross. Four movie watching data runs were collected, each lasting approximately 15 minutes (frames = 921, 918, 915, 901), with subjects passively viewing visual and audio presentations of movie scenes. Movies consisted of both freely available independent films covered by Creative Commons licensing and Hollywood movies prepared for analysis [80]. For both resting state and movie watching data, two runs were acquired with posterior-to-anterior phase encoding direction and two runs were acquired with anterior-to-posterior phase encoding direction.

Structural and functional images were minimally preprocessed according to the description provided in [79]. 7T fMRI images were downloaded after correction and reprocessing announced by the HCP consortium in April, 2018. Briefly, T1w images were aligned to MNI space before undergoing FreeSurfer’s (version 5.3) cortical reconstruction workflow. fMRI images were corrected for gradient distortion, susceptibility distortion, and motion, and then aligned to the corresponding T1w with one spline interpolation step. This volume was further corrected for intensity bias and normalized to a mean of 10000. This volume was then projected to the 2mm *32k_fs_LR* mesh, excluding outliers, and aligned to a common space using a multi-modal surface registration [81]. The resultant cifti file for each HCP subject used in this study followed the file naming pattern: `*_Atlas_MSMAll_hp2000_clean.dtseries.nii`. Resting state and moving watching fMRI images were nuisance regressed in the same manner. Each minimally preprocessed fMRI was linearly detrended, bandpass filtered (0.008-0.25 Hz), confound regressed and standardized using Nilearn’s `signal.clean` function, which removes confounds orthogonally to the temporal filters. The confound regression strategy included six motion estimates, mean signal from a white matter, cerebrospinal fluid, and whole brain mask, derivatives of these previous nine regressors, and squares of these 18 terms. Spike regressors were not applied. Following these preprocessing operations, the mean signal was taken at each time frame for each node, as defined by the Schaefer 400 parcellation [6] in *32k_fs_LR* space.

The movie data included 859 annotations of the movie scenes. These annotations were aligned with TRs so that every volume and every annotation was represented a binary “1” or “0” if that annotation was present or absent. We concatenated these annotations across all four movies and computed a 859 × 859 concordance matrix. We then calculated the mean concordance between all pairs of annotations as 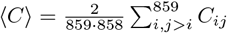. To detect clusters of annotations, we defined the following modularity optimization problem [82]:

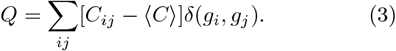

Briefly, *Q* is a quality function that measures the “goodness” of a partition. The summation is over all pairs of nodes *i* and *j*. The Kronecker delta function, *δ*(*x, y*) is 1 when *x* = *y* and 0 otherwise. Effectively, this ensures that the summation is over pairs of nodes that fall within the same community. Intuitively, if the communities are “cohesive” – i.e. *C*_*ij*_ *>>* ⟨*C*⟩ – then high-quality partitions correspond to large values of *Q*.

We used the Louvain algorithm [83] to optimize *Q* with 1000 random restarts. In general, the resulting partitions were not identical. To consolidate these slightly different solutions into a single representative partition, we performed consensus clustering using a previously described approach [84, 85]. This procedure resulted in a single representative partition consisting of 20 annotation clusters. Intuitively, each cluster corresponded to terms that co-occurred with one another above baseline level, where baseline co-occurrence was defined as ⟨*C*⟩.

#### C. Elegans

We analyzed a series of publicly available *C. elegans* recordings made during mating. For details of the recording procedure, please see Susoy *et al*. [48]. Below we paraphrase the description of recordings from Susoy *et al*. [48].

Recordings focused on tail neurons in male *C. elegans* known to be related to mating. Imaging was performed using a custom spinning-disc confocal microscope as described in Venkatachalam *et al*. [86]. Emitted light was split into the red and green channels and captured using two Andor Zyla 4.2 sCMOS cameras. Each camera recorded 256 × 512 pixel area of interest at 200 Hz and with the system pixel size of 0.45 *µ*m. Volumetric imaging was done using a 40×, 1.30 NA Nikon Plan Fluor objective mounted on a piezoelectric stage. Brain volumes were sampled at a rate of 10 Hz, with each volume consisting of 20 optical sections approximately 1.75 *µ*m apart. As described in Susoy *et al*. [48], the male’s tail was tracked by adjusting the microscope stage position with a controller.

Neurons were identified based on a combination of multiple features. These included position, morphology, and expression of fluorescent markers. In total, Susoy *et al*. [48] identified 76 neurons across all datasets.

Neuronal activity traces were extracted from raw image volumes following image pre-processing and registration. All red channel volumes were registered to the green channel through rigid transformation. A difference of Gaussian filter was applied to both channels to suppress the background noise. The datasets were subsampled to include every other volume, which were converted into 5D big-data-viewer arrays and used for neuronal segmentation and tracking with MaMuT 0.27, a Fiji plugin [87, 88]. Segmentation/tracking was carried out quasi-manually, with MaMuT’s automated tracking and a custom neuron tracking tool. Fluorescence intensities (F(red) and F(green)) were extracted by computing mean pixel values for the 2.25 × 2.25 × 3.5 *µ*m volumes surrounding the center of each nucleus in the green and red channels. Savitzky-Golay filtering (polynomial order of 1 and frame length 13) was applied to intensity traces from each channel for noise-reduction [89]. Susoy *et al*. [48] computed neuronal activity as the ratio F(green)/F(red) to minimize the effects of corre-lated noise and motion artifacts. To mitigate remaining motion artifacts, a singular value decomposition was applied to the main datasets retaining 2/3 of components. Besides imaging data, this dataset included a suite of time-varying behavioral measures. Among the measures were:

1. Swimming velocity: centroid velocity of tail neurons AS11, PLM, and EF1.
2. Tail curvature: Angle between rays from neuron PVY and intersecting AS11 and PLM.
3. Distance of the male tail to the tail of the hermaphrodite.
4. Distance of the male tail to the vulva.
5. Spicule protraction: distance between spicule neurons SPD and SPC. During protracting, the distance decreases.

### Edge time series

Generally, FC is defined as statistical dependence of activity recorded at distant sites. Although in principle this definition allows for many different measures of FC [90], in practice it is operationalized as the bivariate product-moment correlation. Most commonly applied to large-scale functional imaging data–e.g. fMRI BOLD–correlation as FC has also been applied to data acquired at single-cell resolution [91] and at much faster temporal resolution [92].

To compute the correlation as FC, suppose we have z-scored two time courses, **z**_*i*_ = [*z*_*i*_(1), …, *z*_*i*_(*T*)] and **z**_*j*_ = [*z*_*j*_(1), …, *z*_*j*_(*T*)]. We calculate their correlation as:

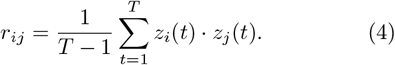

Equivalently, we could define the vector **r**_*ij*_ = [*r*_*ij*_(1), …, *r*_*ij*_(*T*)], where *r*_*ij*_(*t*) = *z*_*i*_(*t*) · *z*_*j*_(*t*). We refer to *r*_*ij*_(*t*) as the instantaneous co-fluctuation between time courses *i* and *j*. Its value is positive if the two time courses are deflecting in the same direction relative to their mean values—i.e. when sign(*z*_*i*_(*t*)) = sign(*z*_*j*_(*t*))– and its amplitude encodes how far from their means the two time series are deflecting in terms of standard deviations squared. Given this time series, we can reinterpret the correlation coefficient as the time-averaged co-fluctuation between any pair of brain regions, *i* and *j*.

### Linear regression models

Here, we fit multilinear models of the form:

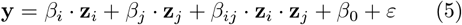

where **y** is the response variable – the behavioral variable – and where **z**_*i*_, **z**_*j*_, and **z**_*i*_**z**_*j*_ are the z-scored activity of node *i*, node *j*, and the edge time series for node pair, {*i, j*}. We estimated the parameters *β*_*i*_, *β*_*j*_, *β*_*ij*_, and *β*_0_ using ordinary least squares. We fit a model of the above form for every pair of nodes, yielding a total of *N* (*N* − 1)*/*2 models.

### Prewhitening

We also considered the effect of prewhitening on our results. Prewhitening is a procedure for reducing the impact of serial correlation in regression analyses. Specifically, prewhitening helps suppress inflated and spurious effects.

Our prewhitening procedure was implemented as follows. First, we estimated regression coefficients as described in the previous section. Using these coefficients, we define the vector **B** = [*β*_*i*_, *β*_*j*_, *β*_*ij*_, *β*_0_] for node pairs {*i, j*}. We also define the matrix **X** = [**z**_*i*_, **z**_*j*_, **z**_*i*_**z**_*j*_, **1**]. The matrices **X** and **B** have dimensions [*T* × 4] and [4 × 1], respectively. We then calculated **Ŷ**= **XB** and calculated the residuals as **E** = **Y** − **Ŷ**.

Then, we fit an autoregressive model of order *p* = 25 to the residuals, **E**. Here, we make one simplifying assumption; rather than estimate autoregressive coefficients separately for each node, we sample 100 random node pairs, obtain autoregressive coefficients separately for each, and take the average. We verified that by 100 samples, the mean values of the coefficients have sufficiently stabilized (adding additional samples does not appreciably change the results).

Next, we compute the prewhitening matrix, **W**. To do this, we estimate the matrix **V**^−1^. We do this efficiently by calculating **AA**^⊺^, where **A** is the banded diagonal matrix with a value of 1 along the main diagonal, the first autoregressive coefficient on the first lower off-diagonal band, the second coefficient on the second band, and so on.

Once we calculate **V**^−1^, we perform singular value decomposition, obtaining eigenvalues, **U** and eigenvectors, **d**. From these outputs, we calculate the prewhitening matrix as **W** = **UD**^1*/*2^**U**^⊺^, where **D** = diag(*d*).

With the prewhitening matrix, we can simply fit the regression model:

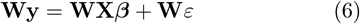

where **X** is the same [*T* × 4] matrix as before and ***β*** is a new set of regression coefficients with dimensions [4 × 1].

## RESOURCE AVAILABILITY

All data are publicly available (zebrafish: https://janelia.figshare.com/articles/dataset/Whole-brain_light-sheet_imaging_data/7272617; *C. elegans*: https://data.mendeley.com/datasets/922xsfxh2g/1; human imaging data are available after signing a data use agreement: https://db.humanconnectome.org/). Code for estimating edge time series is available here: https://github.com/brain-networks/edge-centric_demo. Code for modeling time-varying behaviors as a function of activity and edge time series is available here: https://github.com/brain-networks/ets_glms.

## AUTHOR CONTRIBUTIONS

HM contributed to the conceptualization, investigation, formal analysis, and the writing of the draft along with review and editing of the manuscript. AM contributed to funding acquisition, methodology, and writing of the original draft. RB contributed to conceptualization, data curation, formal analysis, funding acquisition, methodology, visualization, project administration, supervision, and the writing of the draft along with review and editing of the manuscript.

## ACKNOWLEDGEMENTS

RFB and AM acknowledge support from the National Science Foundation (NCS-FO award #2023985).

---

**Figure S1.**
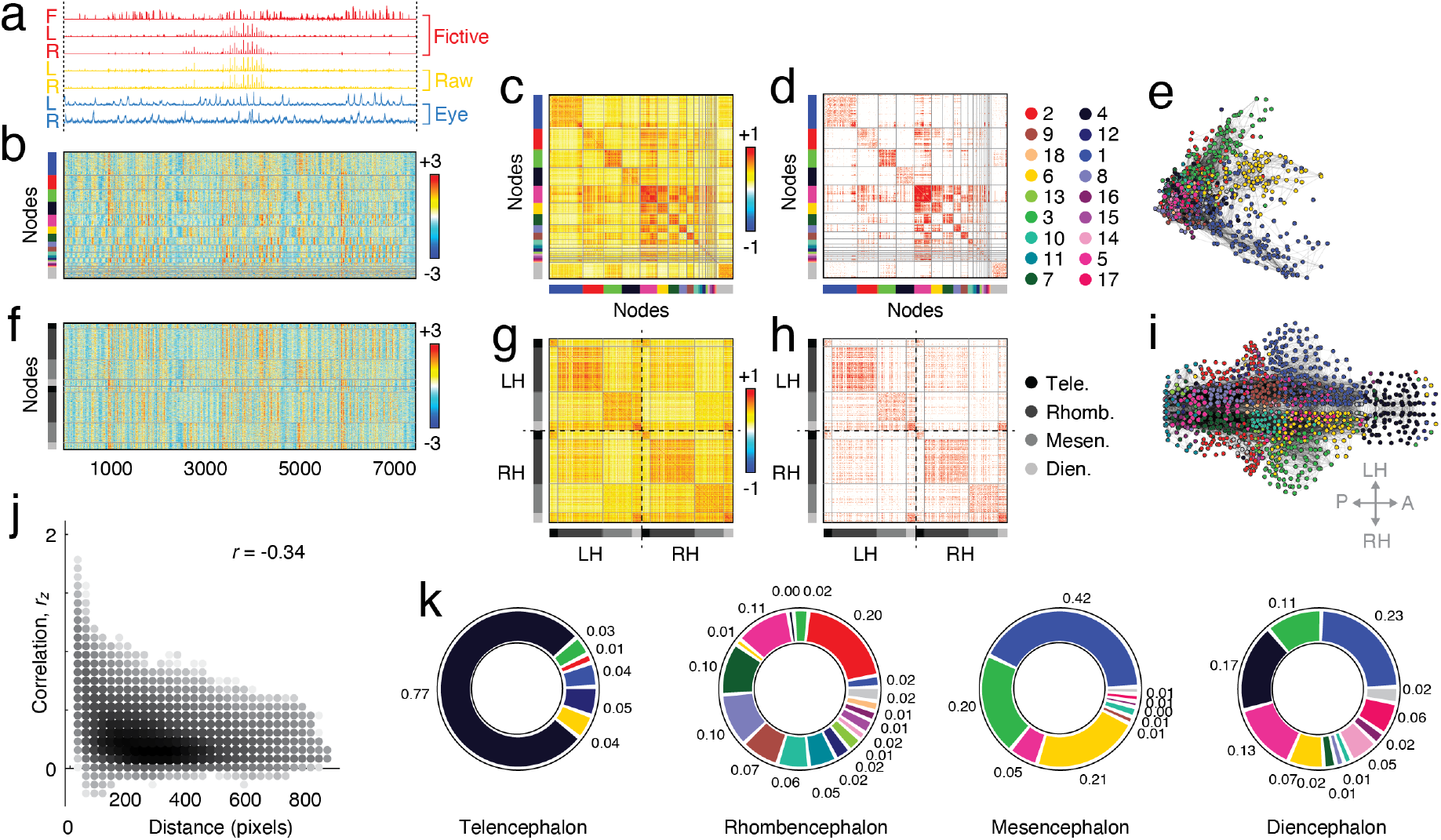
Description of zebrafish connectivity. (*a*) Time-varying measures of inferred motion. (*b*) Z-scored parcel (*N* = 1000) time series ordered by communities. (*c*) Correlation matrix. (*d*) Thresholded correlation matrix. (*e*) Force-directed layout for thresholded matrix. Panels *f* -*h* Show parcel time series, correlation matrix, and thresholded matrix ordered by anatomical label rather than community organization. Panel *i* is analogous to panel *e* but shows network organization in anatomical layout. (*j*) Scatterplot of distance (inter-pixel distance) and correlation magnitude. (*k*) Community composition of four large anatomical areas.

**Figure S2.**
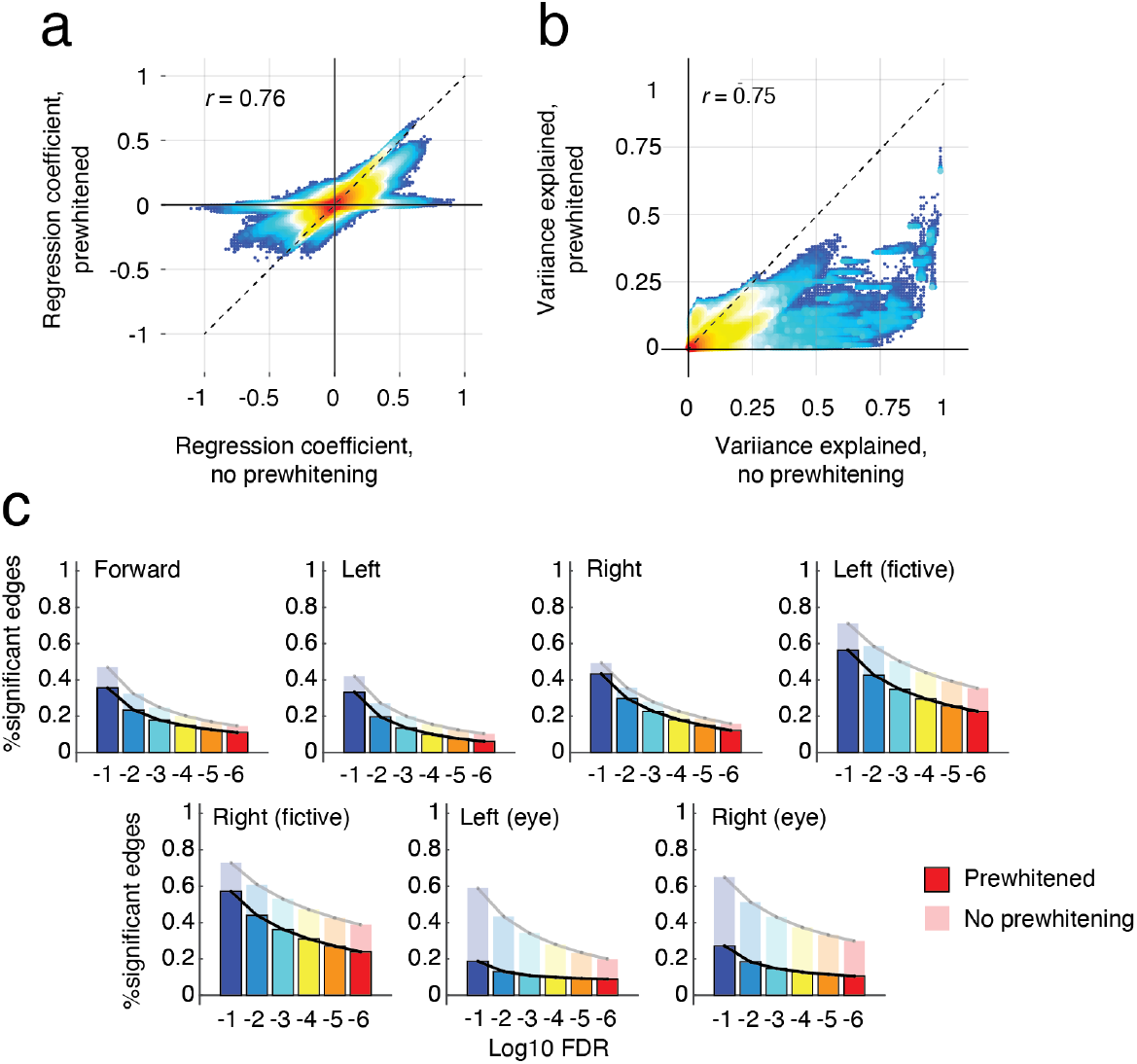
Assessing the impact of pre-whitening on reported results. In the main text we fit linear models to explain time-varying behaviors in terms of activity and connectivity. Importantly, the reported procedure did not include prewhitening–a processing step that reduces the impact of serial correlation, which can artifactually inflate effect sizes. Here, we compare parameters of models fit both with and without any prewhitening. (*a*) Two-dimensional histogram of regression coefficients. Here, coefficients are aggregated across all 11 fish, 7 behaviors, and all edges. (*b*) Two-dimensional histogram of variance explained. Note that, in general, variance explained is reduced for the prewhitened model. Panel *c* depicts fraction of significant edges for each of the seven behaviors. The semi-transparent bar plots depict models results from models fit without prewhitening; the opaque bar plots show results from models fit following prewhitening. Note, again, that the prewhitening procedure reduces the number of significant edges in all cases.

**Figure S3.**
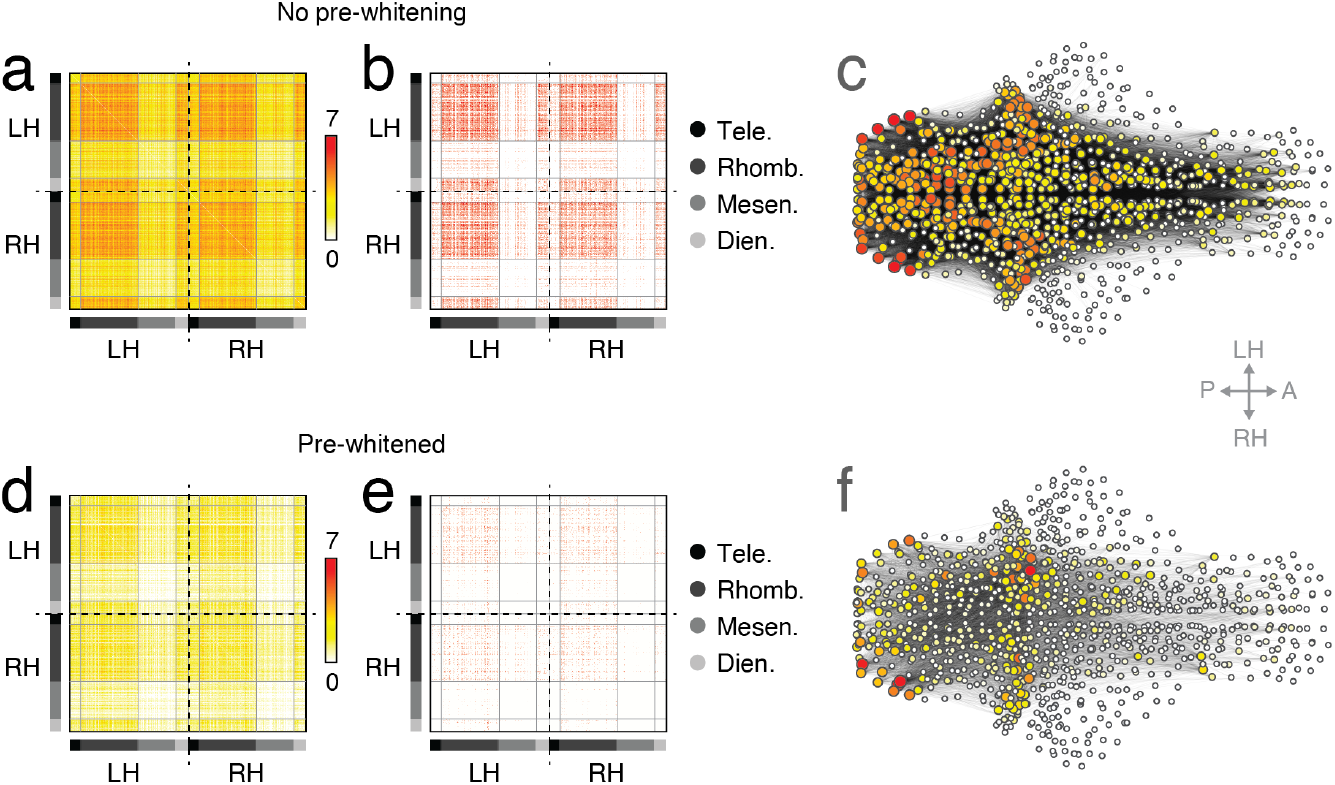
Overlap of statistically significant edges. (*a*) Mean number of behaviors for which a given edge (interaction) was statistically significant. Panel *b* shows edges with count equal to 7 (all behaviors) for the non-pre-whitened data. (*c*) Row/column sum of the entries in panel *b* projected back into anatomical space. Warmer (redder) colors correspond to nodes whose edges tended to be statistically significant. Panels *d* -*f* show analogous figures but for data that had undergone pre-whitening.

**Figure S4.**
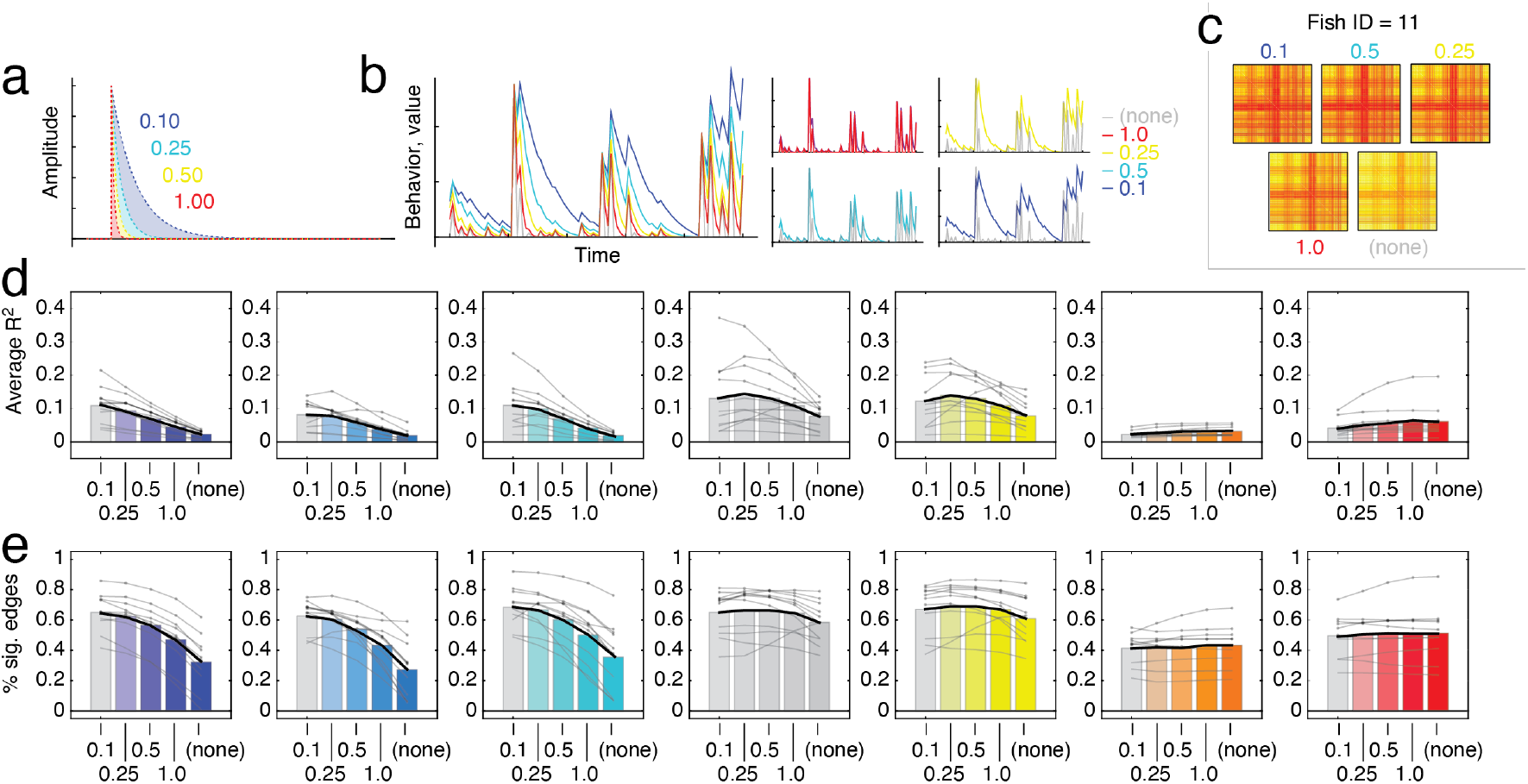
Convolution of behavioral data with temporal kernel. Here, we convolve zebrafish behavioral data with four different kernels corresponding to decaying exponential functions with different rate parameters (see panel *a* for shape of each kernel and panel *b* for effect of each kernel of small amount of behavioral data from a representative fish. In general the decision to smooth or not impacts model fit. In panel *c* we show an example matrix of *R*^2^ value for a representative fish for forward movement. Note that the intensity of entries varies as a function of rate parameter; sharper decay (or no convolution) lead to the smallest *R*^2^ values, generally. We show this in panel *d* for each of the seven behavioral measures. For kinematic measures (forward/left/right movement), *R*^2^ increases more or less monotonically as the slope of the exponential grows shallower. For eye movements, *R*^2^ changes very little. We observe a corresponding effect when we calculate the fraction of edges that pass tests for statistical significance (*q* = 0.01). In general, convolution either increases the number of statistically significant edges or it leaves that number relatively unchanged.

**Figure S5.**
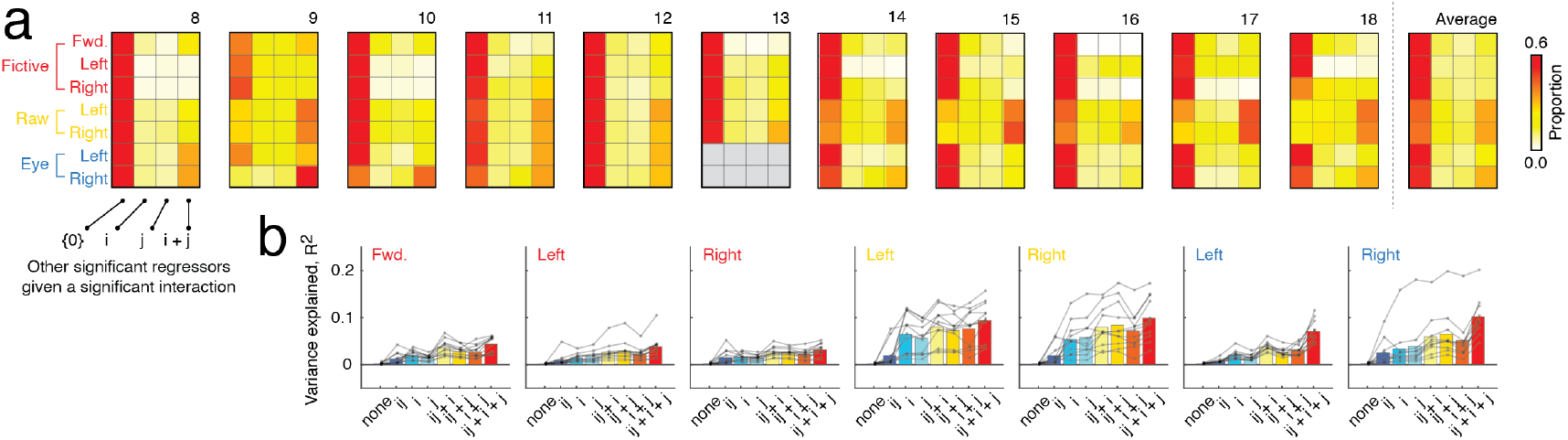
Further characterization of statistically significant interactions. First, we identified all pairs, {*i, j*}, with statistically significant interactions (with false discovery rate fixed at *q* = 0.01). For these pairs, we then examined whether activations *i, j*, or *i* + *j* were also signification. (*a*) In general, we find that most statistically significant interactions occur when activations are not statistically significant. Next, we labeled all node pairs, {*i, j*}, based on which terms were statistically significant. For each label, we then calculated the mean variance explained. Panel *b* shows the results of this exercise. In general, models for which all three regressors were statistically significant were associated with greater *R*^2^ than other models.

**Figure S6.**
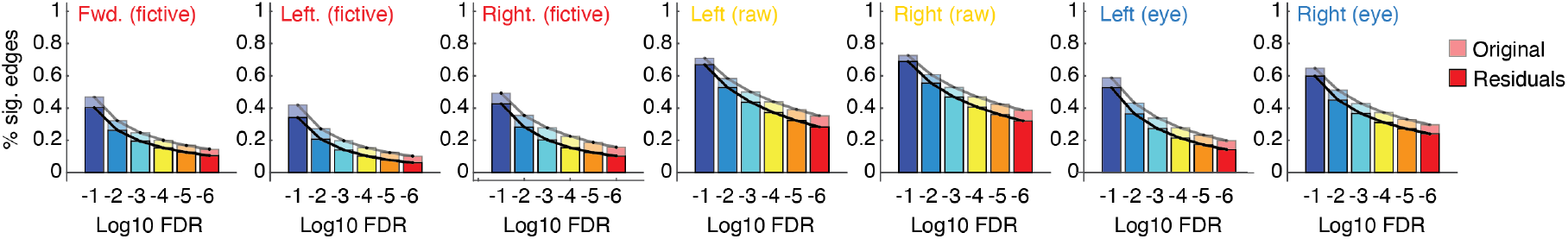
Analysis of residualized behavior. In the main text, we fit models of the form **y** = *β*_*i*_**z**_*i*_ + *β*_*j*_ **z**_*j*_ + *β*_*ij*_ **z**_*i*_ · **z**_*j*_ + *β*_0_ + *ε*. These models allow edge time series and activations to compete in explaining behavior, **y**. A more conservative version of is to fit the model: **y** = *β*_*i*_**z**_*i*_ + *β*_*j*_ **z**_*j*_ + *β*_0_ + *ε* and use its parameters to obtain the estimate **ŷ** and calculate the residuals, **ŷ** − **y**. This allows the two activation time courses, **z**_*i*_ and **z**_*j*_, to explain as much variance as possible without any competition against the interaction term, **z**_*i*_ · **z**_*j*_. We could then fit the model: **ŷ** − **y** = *β*_*ij*_ **z**_*i*_ · **z**_*j*_ + *β*_0_ + *ε*. Here, we report the results of this model, focusing on the fraction of edges that are statistically significant. In general, we find that residualized models exhibit fewer significant edges (on average) a 24.4 *±* 18.6% reduction. Nonetheless, fewer than 1% (0.67%) of models exhibited no statistically significant edges. form **y** = *β*_*i*_**z**_*i*_ + *β*_*j*_ **z**_*j*_ + *β*_*ij*_ **z**_*i*_ · **z**_*j*_ + *β*_0_ + *ε*. These models allow edge time series and activations to compete in explaining

**Figure S7.**
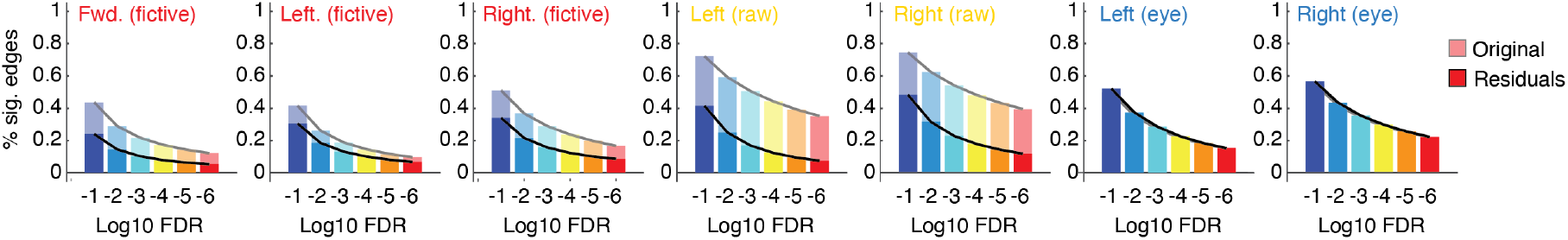
Effect of including principal components in regression models. In the main text, we fit models of the *β*_*k*_*ϕ*_*k*_ + *β*_0_ + *ε*. Here, *ϕ*_*k*_ is the *k*th principal component of the complete set of neural time courses. This model allows for activity beyond that of nodes *i* and behavior, **y**. Here, we fit models of the form: 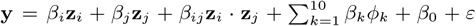. Here, *ϕ*_*k*_ is the *k*th *j* to account for variance that could otherwise be explained by edge, *i, j*. Again, this approach is more conservative in that it reduces the fraction of significant edges, but because we find robust evidence of statistically significant interactions, it is generally in line with the hypothesis that edges carry explanatory power.

**Figure S8.**
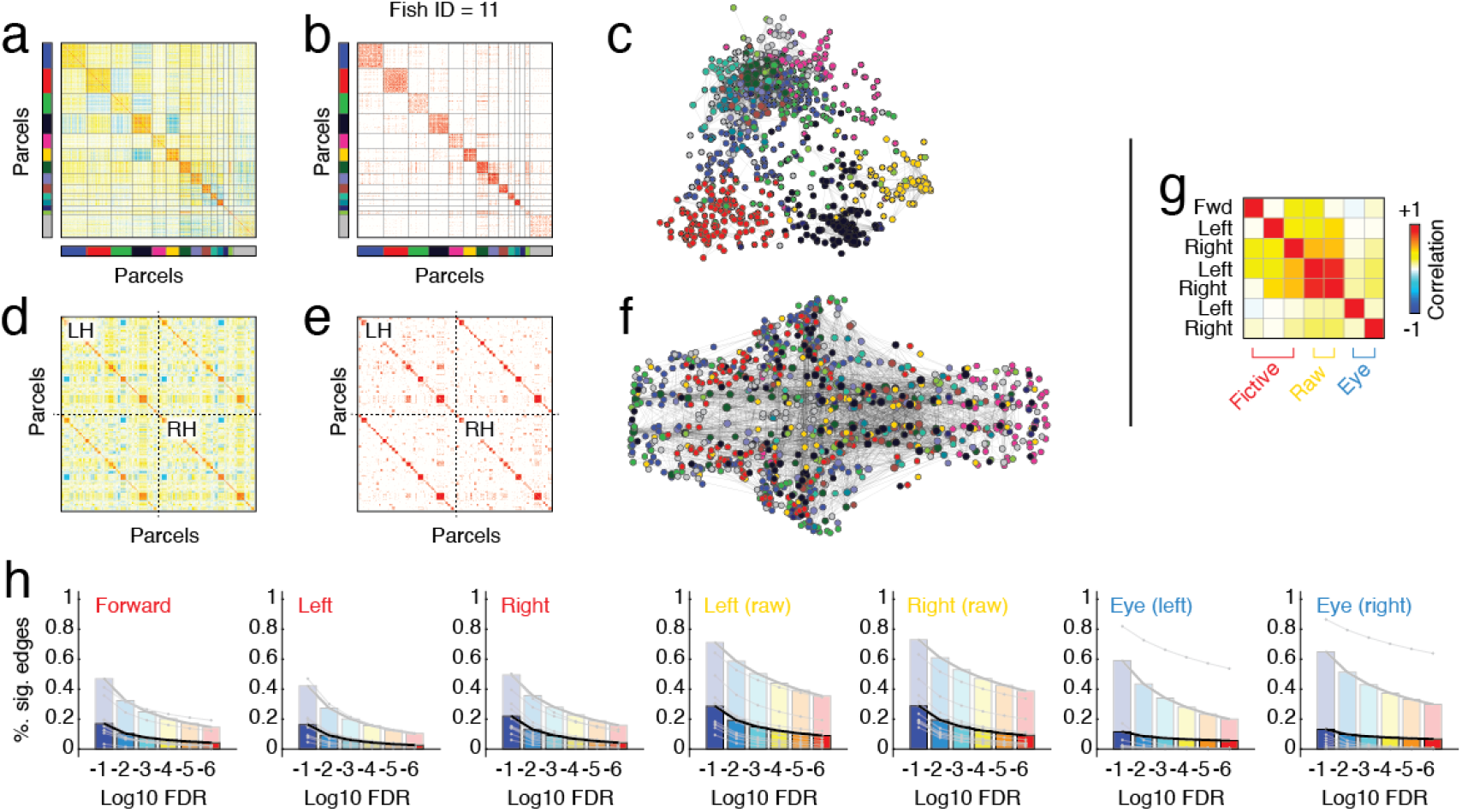
Effect of individualized and functional parcellations. In the main text, we used a division of neurons into a set of group-level parcels that were shared across individuals. Here, we show results using fish-specific and functionally-defined parcels. Specifically, we started with a coarse functional parcellation (clusters defined based on neuron-to-neuron correlation structure alone), and subdivided those clusters using neurons’ 3D coordinates (mirrored into a single hemisphere) into spatially compact sub-clusters. In panels *a*-*f* we show raw and thresholded correlation structure sorted by module (*a* and *b*) and by hemisphere (*d* and *e*). Panels *e* and *f* depict the thresholded network with coordinates defined based on force-directed layout and anatomy. We fit edge-level regression models independently for each fish using pre-whitened data. We then calculated the similarity of regression coefficients across all seven measures (see panel *g*, which averages these similarity values across fish). (*h*) Fraction of statistically significant edges at varying statistical thresholds. The semi-transparent plots show data using group parcels.

**Figure S9.**
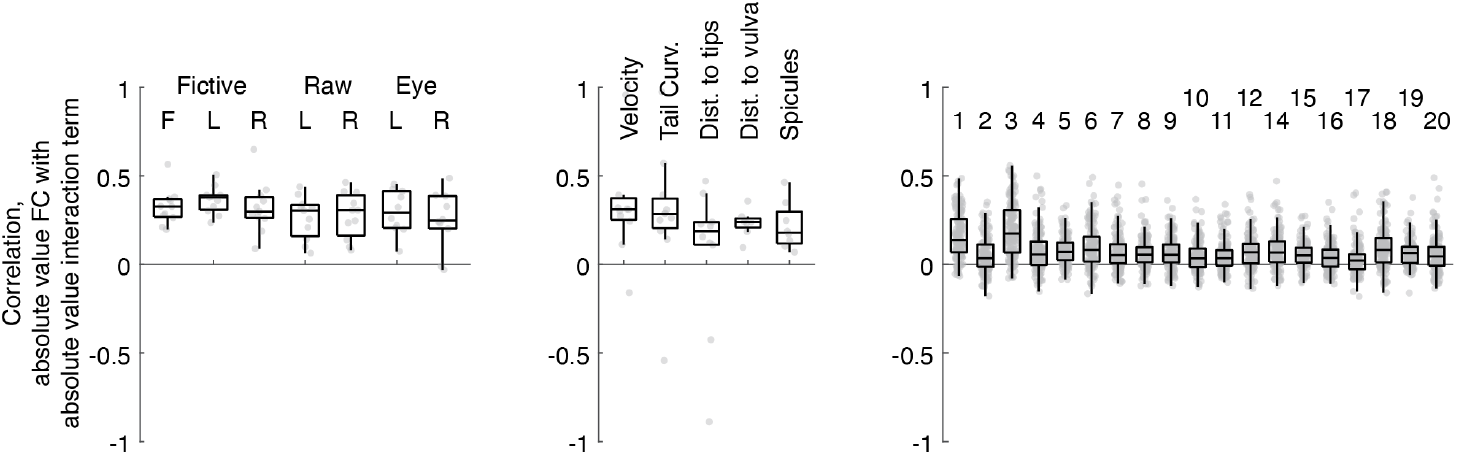
Linking FC with *β*_*ij*_. Here, we show boxplots for each behavioral term and all three organisms wherein we calculate the correlation between the absolute weight of FC with the absolute value of the interaction term, *β*_*ij*_. In general, we find a positive relationship that holds across brains and behavioral measures.

